# Epigenetically regulated p53 activity maintains intestinal regulatory T cell identity to prevent inflammation

**DOI:** 10.1101/2025.08.29.673118

**Authors:** Stephanie Silveria, Janneke GC Peeters, Jenna Vickery, Giovana MB Veronezi, Srinivas Ramachandran, Michel DuPage

## Abstract

Regulatory T cells (Tregs) are critical guardians of immune homeostasis that must operate in diverse and often inflammatory conditions. However, the mechanisms that Tregs use to maintain their stability and function, especially in response to the stresses of distinct microenvironments, remain incompletely understood. Previous work identified the repressive chromatin modification histone 3 lysine 27 trimethylation (H3K27me3) as a rheostat for Treg function. Here, we find that loss of H3K27me3 in Tregs activates the tumor suppressor p53. Stabilization of p53 using the MDM2 inhibitor Nutlin-3 protected Tregs from losing their master transcription factor Foxp3 *in vitro* when cultured with the Th17 cytokines IL-6 and IL-1β, while p53 deficiency rendered Tregs more prone to Foxp3 loss. Treg-specific p53 deficiency resulted in accumulation of cells that had lost Foxp3 expression (“ex-Tregs”) and reduction of suppressive markers on Tregs specifically in the colon. Additionally, these mice exhibited inflammation in the colon at homeostasis and increased severity of induced colitis. These results demonstrate a specific role for p53 in the maintenance of Treg stability in Th17-polarizing environments and present a possible target for improving Treg-based immunotherapies for diseases defined by intestinal inflammation, such as inflammatory bowel disease (IBD).

## Introduction

Regulatory T cells (Tregs) are an essential immunosuppressive subset of CD4^+^ T cells responsible for preventing autoimmunity, controlling inflammation, and promoting tissue repair^1–3^. Epigenetic modifications in Tregs, including direct regulation of the Treg master transcription factor locus *Foxp3*, are critical for Treg suppressive function and maintenance of their lineage identity^4–6^. At the *Foxp3* locus, DNA hypomethlyation of the conserved non-coding sequence 2 (CNS2) region, known as Treg-specific hypomethylated region (TSDR), results in recruitment of transcription factors that help to stabilize Foxp3 expression in Tregs^4,7–15^. Chromatin-modifying enzymes have also been shown to contribute to Treg identity through repression of pro-inflammatory genes^16–18^. The methyltransferase Enhancer of Zeste Homolog 2 (EZH2), a component of the polycomb repressive complex 2 (PRC2), promotes Foxp3’s repressive transcriptional program through deposition of H3K27me3 after activation^16^. Genetic deletion of *Ezh2* in Tregs severely compromises their stability and function, resulting in increased generation of ex-Tregs and impaired immune homeostasis^19–22^. In contrast, hyperactivation of EZH2 promoted Treg stability and poised the cells for differentiation, improving their ability to suppress autoimmunity^23^. Additionally, while mice harboring *Ezh2*-deficient Tregs displayed enhanced anti-tumor immune responses, deficiency for the lysine demethylase *Jmjd3* in Tregs resulted in accelerated tumor growth, further suggesting that H3K27me3 may act as a switch for Treg function^24^. Despite the importance of proper H3K27me3 distribution upon activation, the pathways that are controlled by this modification and their roles in Treg function have not been extensively investigated.

The tumor suppressor p53, encoded by *Trp53* in mice, is a transcription factor that controls a vast network of cellular responses to stress. While best known for its roles in apoptosis and cell cycle arrest in response to DNA damage, it is also involved in more diverse biological processes such as cellular metabolism, autophagy, and redox biology^25–28^. In the immune system, p53 has been implicated in immune regulation and tolerance^29–35^. p53-deficient mice suffer more severe disease in the autoimmune models collagen-induced arthritis (CIA), antigen-induced arthritis, experimental autoimmune encephalomyelitis (EAE), streptozotocin (STZ)-induced diabetes and type 1 diabetes (T1D), emphasizing the protective role that p53 plays in the context of autoimmunity^36–39^. Mice with T cell-specific ablation of p53 spontaneously developed autoimmune pathology as they aged, demonstrating that p53 in T cells is essential for preventing autoimmunity^40^. Treg differentiation from p53-deficient CD4^+^ T cells was also reported to be defective^40,41^. However, whether p53 deficiency affects the maintenance of Treg lineage stability and whether Treg-specific p53 deficiency alone promotes autoimmune disease has not yet been explored.

Tregs play a vital role in maintaining immune homeostasis in the intestinal mucosa, where tolerance to commensal bacteria and dietary antigens is crucial. Disruption of the balance between Th17 effector T cells and Tregs can result in inflammation that contributes to the development of inflammatory bowel disease (IBD), which includes ulcerative colitis and Crohn’s disease^42–44^. Treg dysfunction has been implicated in intestinal inflammation and IBD pathology, and Treg-targeted approaches have been proposed as potential therapeutic interventions for IBD^42,45–51^. Gaining a greater understanding of the unique mechanisms employed by intestinal Tregs to maintain their stability and function in the complex gut microenvironment may contribute to improved therapies.

In this study, we show that global reductions in the levels of H3K27me3 modifications in Tregs activates the p53 pathway. Increased p53 activity promotes the maintenance of Foxp3 expression in Tregs *in vitro* when cultured with the Th17 cytokines IL-1β and IL-6 and *in vivo* in the colon. Interestingly, the apparent increased maintenance of Foxp3 expression in Tregs was not due to the increased apoptosis or cell cycle arrest of Tregs that lost Foxp3. Specific deficiency for p53 in Tregs led to more ex-Tregs and a reduction of suppressive markers on Tregs in the colon. Furthermore, Treg-specific p53-deficiency led to increased inflammation in the colon of mice at steady-state and increased the severity in models of induced colitis. Thus, while not essential for global Treg stability, p53 is necessary to maintain Treg cell identity in the Th17-polarizing environment of the colon.

## Results

### Regulation of H3K27me3 controls Foxp3 maintenance in Tregs

We used Foxp3 lineage tracing (*Foxp3-GFP-hCre;R26^LSL-RFP^*) mice to assess Foxp3 maintenance in regulatory T cells (Tregs) with genetically altered levels of H3K27me3 *in vivo* (Figure 1A). We observed that splenic *Ezh2*-deficient (*Ezh2^Δ/Δ^)* Tregs were more prone to losing Foxp3 expression, generating more ex-Tregs, whereas *Jmjd3*-deficient (*Jmjd3^Δ/Δ^*) Tregs better maintained Foxp3 expression, generating fewer ex-Tregs, when compared to wildtype (*WT*) Tregs (Figures 1B-D). To further investigate the role of H3K27me3 in Treg reprogramming, we also tested Foxp3 stability *in vitro* with each Treg genotype after anti-CD3/CD28 stimulation and IL-2 (Figure 1E)^52^. However, under these conditions *in vitro*, and unlike *in vivo*, all Tregs equally maintained Foxp3 and did not generate ex-Tregs (Figure 1F, top row). We hypothesized that these *in vitro* conditions must not fully recapitulate the destabilizing conditions encountered by Tregs of these genotypes *in vivo*. Therefore, we tested whether the addition of pro-inflammatory cytokines to the Th0 (IL-2 only) condition would better mimic the conditions encountered *in vivo* that destabilized Foxp3 expression by using conditions for the differentiation of Th1 (IL-2+IL-12), Th2 (IL-2+IL-4), or Th17 (IL-2+IL-1β+IL-6+IL-23) cells (Figure 1E)^53–58^. Analysis after varying the cytokines treated in culture demonstrated that *Ezh2^Δ/Δ^*Tregs in Th17 conditions lost more Foxp3 expression compared to *WT* Tregs, while *Jmjd3^Δ/Δ^* Tregs lost less (Figures 1F-H and Extended Data Figures 1A-B). In addition, while Th2 or Th1 conditions did not promote significant changes in Foxp3 expression in either genotype, Th1 conditions increased IFN-γ production in *Ezh2^Δ/Δ^* Tregs and decreased IFN-γ production in *Jmjd3^Δ/Δ^* Tregs compared to *WT* (Extended Data Figures 1C-D)^20,22,59^. Thus, reprogramming of Tregs *in vitro* necessitates the addition of pro-inflammatory cytokines to recapitulate aspects of Treg reprogramming observed *in vivo*.

**Figure 1.**
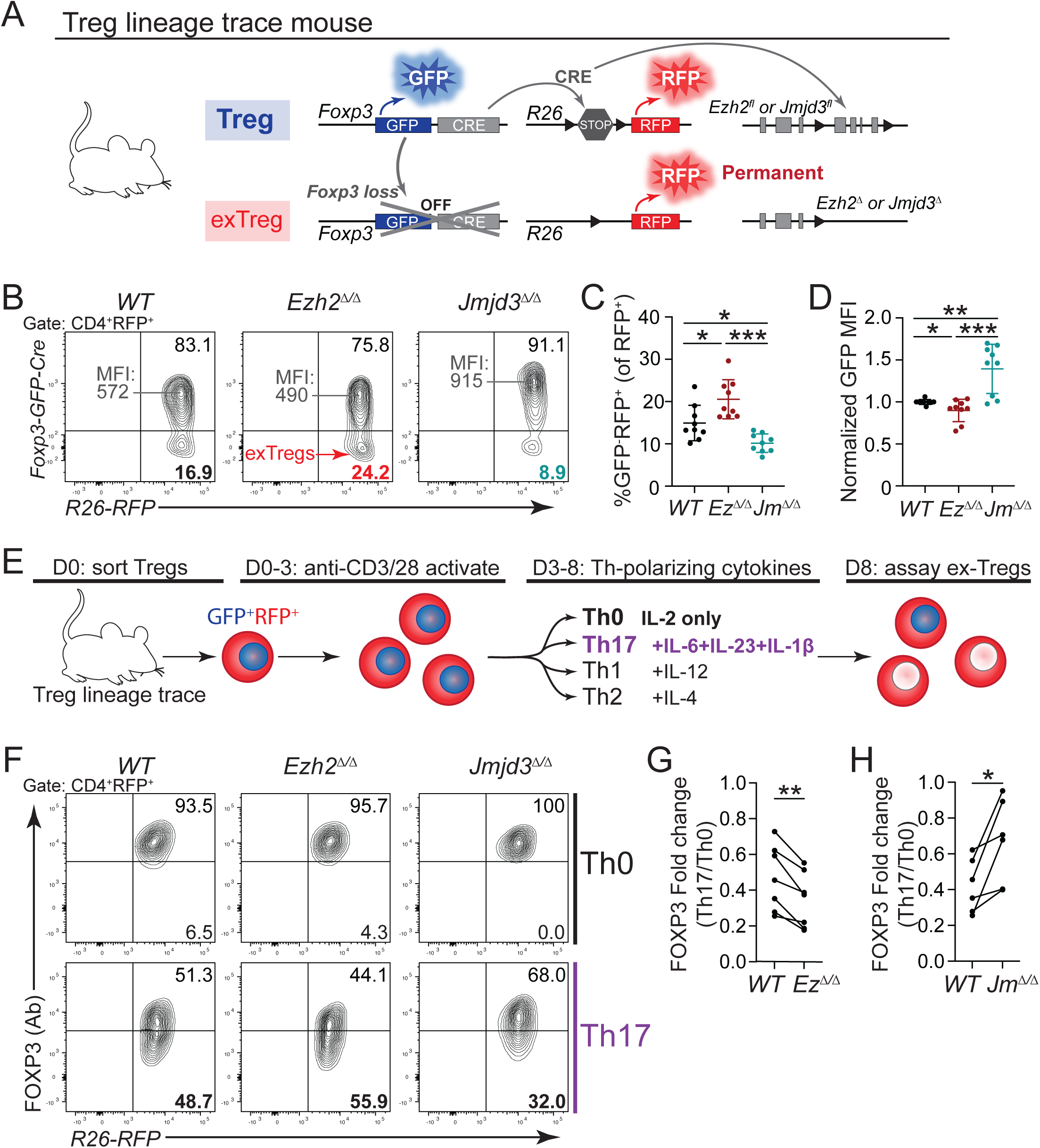
Regulation of H3K27me3 controls Foxp3 maintenance in Tregs. (A) Schematic of Treg lineage trace system. (B) Representative flow plots and frequencies of GFP-RFP+ ex-Tregs and representative GFP MFI of GFP+RFP+ Tregs in splenic CD4+RFP+ cells from WT, Treg.Ezh2 Δ/Δ (EzΔ/Δ), or Treg.Jmjd3 Δ/ Δ (Jm Δ/Δ) mice. (C) Frequency of splenic GFP-RFP+ ex-Treg of total RFP+ cells from WT, Treg.Ezh2 Δ/Δ (EzΔ/Δ), or Treg.Jmjd3 Δ/Δ (Jm Δ/Δ) mice. Data from n=9 mice per genotype. (D) MFI of GFP expression in splenic GFP+RFP+ Tregs from WT, Treg.Ezh2 Δ/Δ (EzΔ/Δ), or Treg.Jmjd3 Δ/Δ (Jm Δ/Δ) mice. (E) Assay design for in vitro Treg reprogramming. (F) Representative flow plots of sorted Tregs in Th0 (top) or Th17 (bottom) conditions on day 8. (G-H) Paired quantification of FOXP3 fold change in Th17 relative to Th0 conditions for WT versus (G) Ez Δ/Δ or (H) Jm Δ/ Δ Tregs. Data points represent biological replicates pooled from six to seven independent experiments. *p<0.05, **p<0.01, ***p<0.001, ****p<0.0001 by Ordinary one-way ANOVA (C), unpaired Student’s t-test (D), or paired Student’s t-test (G-H), mean ± s.d.

### Altered H3K27me3 levels control the activation of the p53 pathway in Tregs

To explore transcriptional changes that might account for the observed differences in Foxp3 maintenance in *WT* versus *Ezh2^Δ/Δ^*or *Jmjd3^Δ/Δ^* Tregs, we performed RNA-seq before and after the *in vitro* activation of FACS-purified naïve thymically-derived Tregs from the different genetic backgrounds^24^. Ingenuity Pathway Analysis (IPA) of differentially expressed genes between *WT* and each mutant Treg genotype revealed an enrichment in genes that are regulated by the transcription factor *Trp53*, or p53 (Figure 2A). *Ezh2^Δ/Δ^* Tregs had a positive z-score for this group of genes, while *Jmjd3^Δ/Δ^*Tregs had a negative z-score, indicating that p53 target genes were more likely to be activated in *Ezh2^Δ/Δ^* Tregs and inhibited in *Jmjd3^Δ/Δ^*Tregs compared to *WT* Tregs. Among all the p53-regulated genes that were changed in an opposing fashion between *Ezh2^Δ/Δ^* and *Jmjd3^Δ/Δ^*Tregs, more than 80% of these genes were upregulated in *Ezh2^Δ/Δ^*, and downregulated in *Jmjd3^Δ/Δ^*, Tregs (Figure 2B). Using published meta-analysis of datasets and single-gene studies assessing direct p53 binding across the genome^60,61^, we identified genes from our IPA analysis in Figure 2B that were confirmed to be directly bound by p53 (Figure 2C). These 11 genes represent direct p53 target loci that were regulated in opposing fashion in *Ezh2^Δ/Δ^* and *Jmjd3^Δ/Δ^* Tregs, 10 of which were upregulated in *Ezh2^Δ/Δ^*Tregs. Next, we confirmed that these genes identified by RNA-seq were induced in *Ezh2^Δ/Δ^* Tregs compared to *WT* Tregs by quantitative PCR (Figure 2D and Extended Data Figure 2A). Furthermore, we verified that their induction was p53-dependent by showing that their induced expression in *Ezh2^Δ/Δ^*Tregs was prevented by *p53*-deficiency in *Ezh2^Δ/Δ^*;*p53^Δ/Δ^*Tregs (Figure 2D and Extended Data Figure 2A). We also found that the canonical p53 target genes *Cdkn1a* and *Bbc3* were upregulated in *Ezh2^Δ/Δ^* Tregs in a p53-dependent manner by qPCR analysis (Extended Data Figure 2B). These genes trended towards upregulation but did not reach statistical significance in the RNA-seq dataset. Expression of the *Trp53* RNA message was not upregulated in *Ezh2^Δ/Δ^* Tregs (Extended Data Figure 2C). However, consistent with p53-dependent increases in p53 target gene expression, the p53 protein was detected in *Ezh2^Δ/Δ^* Tregs compared to *WT* and *Jmjd3^Δ/Δ^* Tregs or conventional CD4^+^ T cells (Tconv) by both western blotting and flow cytometric analyses (Figures 2E and Extended Data Figure 2D). Together, these data demonstrate an increase in p53 protein and p53 transcriptional activity in *Ezh2^Δ/Δ^* compared to *WT* Tregs.

**Figure 2.**
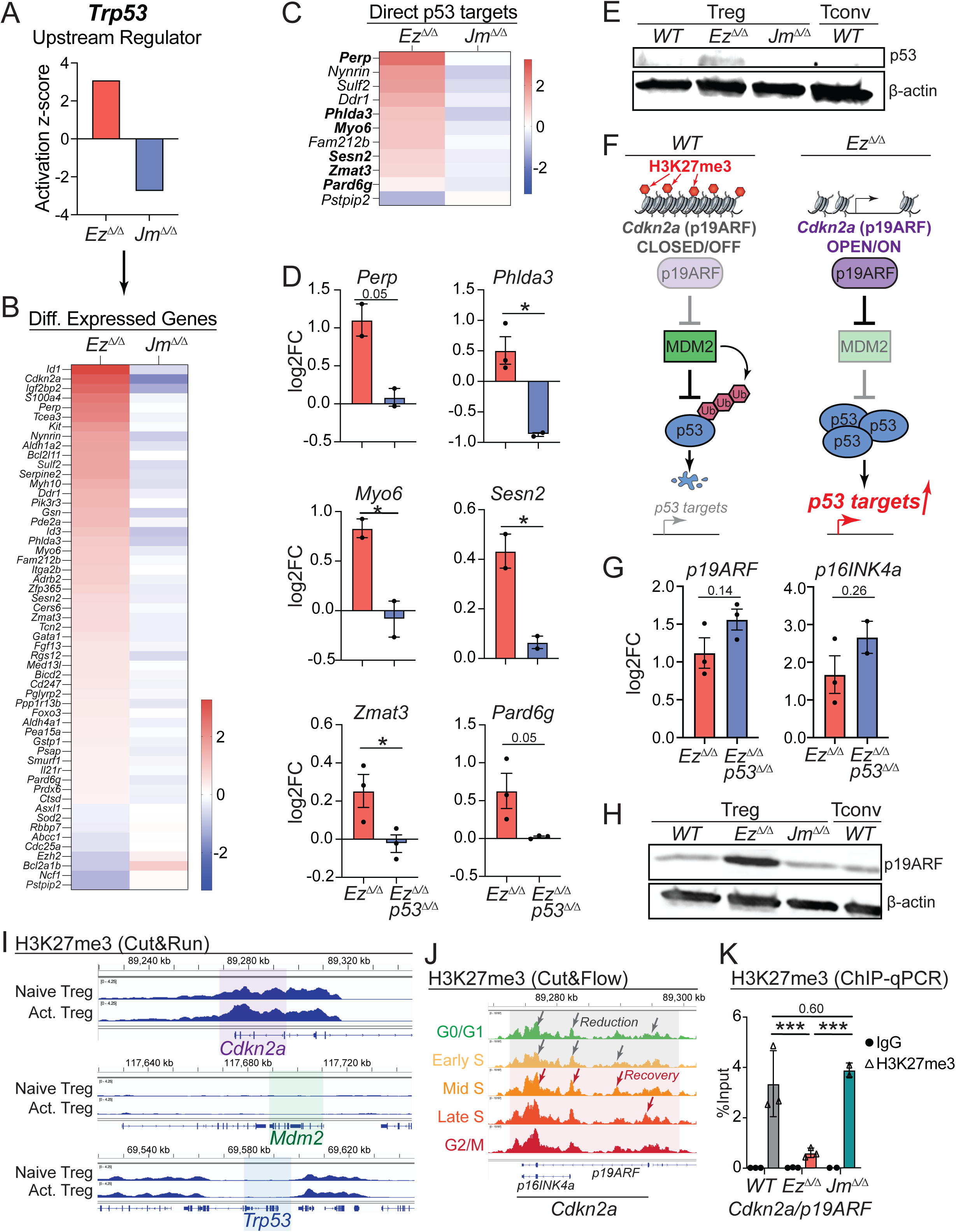
Altered H3K27me3 levels control the activation of the p53 pathway in Tregs. (A) Ingenuity Pathway Analysis activation z-score for Trp53/TP53 as an Upstream Regulator for Ezh2 Δ/Δ vs. WT (Ez Δ/Δ) and Jmjd3 Δ/Δ vs. WT (Jm Δ/Δ) datasets. (B) Heat map of genes identified from the Ingenuity Pathway Analysis Trp53/TP53 Upstream Regulator list that were differentially expressed between Ezh2 Δ/Δ vs. WT (Ez Δ/Δ) and Jmjd3 Δ/Δ vs. WT (Jm Δ/Δ) datasets. (C) Heat map of genes from (B) identified as confirmed direct p53 targets (Fischer 2017). (D) qPCR log2 fold change analysis of select p53 targets in Ezh2 Δ/Δ (EzΔ/Δ) or Ezh2 Δ/ Δp53Δ/Δ (EzΔ/Δp53Δ/Δ) versus WT Tregs activated 4 days in vitro. Data points represent biological replicates pooled from two to three independent experiments. (E) Western blot of p53 protein in WT, Ez Δ/Δ, or Jm Δ/Δ Tregs or WT conventional CD4+ T cells (Tconv) activated 4 days in vitro. (F) Schematic of Cdkn2a/p19ARF and p53 activation in WT versus Ez Δ/Δ Tregs. (G) qPCR log2 fold change analysis of EzΔ/Δ or EzΔ/Δp53Δ/Δ versus WT Tregs activated 4 days in vitro. Data points represent biological replicates pooled from two to three independent experiments. (H) Western blot of p19ARF protein in WT, Ez Δ/Δ, or Jm Δ/Δ Tregs or WT conventional CD4+ T cells (Tconv) activated 4 days in vitro. (I) H3K27me3 CUT&RUN analysis of Naïve versus 4 day in vitro-activated (Act.) Tregs for the indicated loci. Data pooled from n=3 mice. (J) H3K27me3 cUt&FLOW analysis of Cdkn2a locus in Tregs activated 4 days in vitro. Representative of n=2 mice. (K) H3K27me3 ChIP-qPCR analysis of Cdkn2a/p19ARF locus in 4 day in vitro-activated WT, EzΔ/Δ, and Jm Δ/Δ Tregs. Data points represent biological replicates. For all plots, *p<0.05, **p<0.01, ***p<0.001, ****p<0.0001 by unpaired Student’s t-test, mean ± s.d.

p53 protein levels are regulated post-transcriptionally by polyubiquitination and proteasomal degradation^62,63^. A potential source of p53 protein stabilization is increased expression of *Cdkn2a*, which encodes the tumor suppressor p19ARF that negatively regulates MDM2, an E3 ubiquitin ligase for p53 (Figure 2F). RNA-seq showed that *Cdkn2a* was among the most upregulated genes compared to *WT* in *Ezh2^Δ/Δ^* Tregs and most downregulated genes in *Jmjd3^Δ/Δ^* Tregs (Figure 2B). Two transcripts are made from *Cdkn2a* – *p19ARF* and *p16ink4a*, and both transcripts were increased equivalently in *Ezh2^Δ/Δ^* Tregs and *Ezh2^Δ/Δ^;p53^Δ/Δ^* Tregs (Figure 2G). This demonstrates that the upregulation of *Cdkn2a* RNA message with *Ezh2*-deficiency was p53-independent. p19ARF protein levels were similarly increased by western blotting (Figure 2H). In line with H3K27me3 directly regulating *Cdkn2a* expression, CUT&RUN analysis showed significant levels of H3K27me3 at the *Cdkn2a* locus in Tregs, which increased upon activation (Figure 2I). However, H3K27me3 modifications were not present at the *Mdm2* or *Trp53* loci (Figure 2I). Using CUT&FLOW analysis of H3K27me3 in Tregs, we found that H3K27me3 increased dynamically at the *Cdkn2a* locus during Treg activation (Figure 2J)^64^. Finally, ChIP-qPCR analysis of H3K27me3 at the *p19ARF* locus confirmed that *in vitro*-activated *Ezh2^Δ/Δ^* Tregs had reduced H3K27me3 levels compared to *WT* (Figure 2K). Overall, these data support the hypothesis that *Cdkn2a/p19ARF* may serve as a sentinel locus to sense the loss of H3K27me3 in cells, triggering p53 stabilization and transcriptional activity at target genes in response to global reduction of H3K27me3 levels.

### p53 activity promotes the maintenance of Foxp3 expression

To test whether p53 activity is important for the maintenance of Foxp3 expression, we compared *WT* and *p53*-deficient (*Foxp3-GFP-hCre;R26^LSL-RFP^;p53^fl/fl^ = p53^Δ/Δ^*) Tregs activated and cultured *in vitro* in Th0 or Th17 conditions, which exhibited the greatest amount of Foxp3 loss in Tregs (Figure 1E). As observed with *Ezh2^Δ/Δ^*compared to *WT* Tregs, *p53^Δ/Δ^* Tregs maintained Foxp3 expression normally in Th0 conditions (Figure 3A and Extended Data Figure 3A). However, *p53^Δ/Δ^* Tregs lost significantly more Foxp3 when cultured in Th17 conditions, suggesting that p53 function promotes the maintenance of Foxp3 expression in Tregs (Figure 3A).

**Figure 3.**
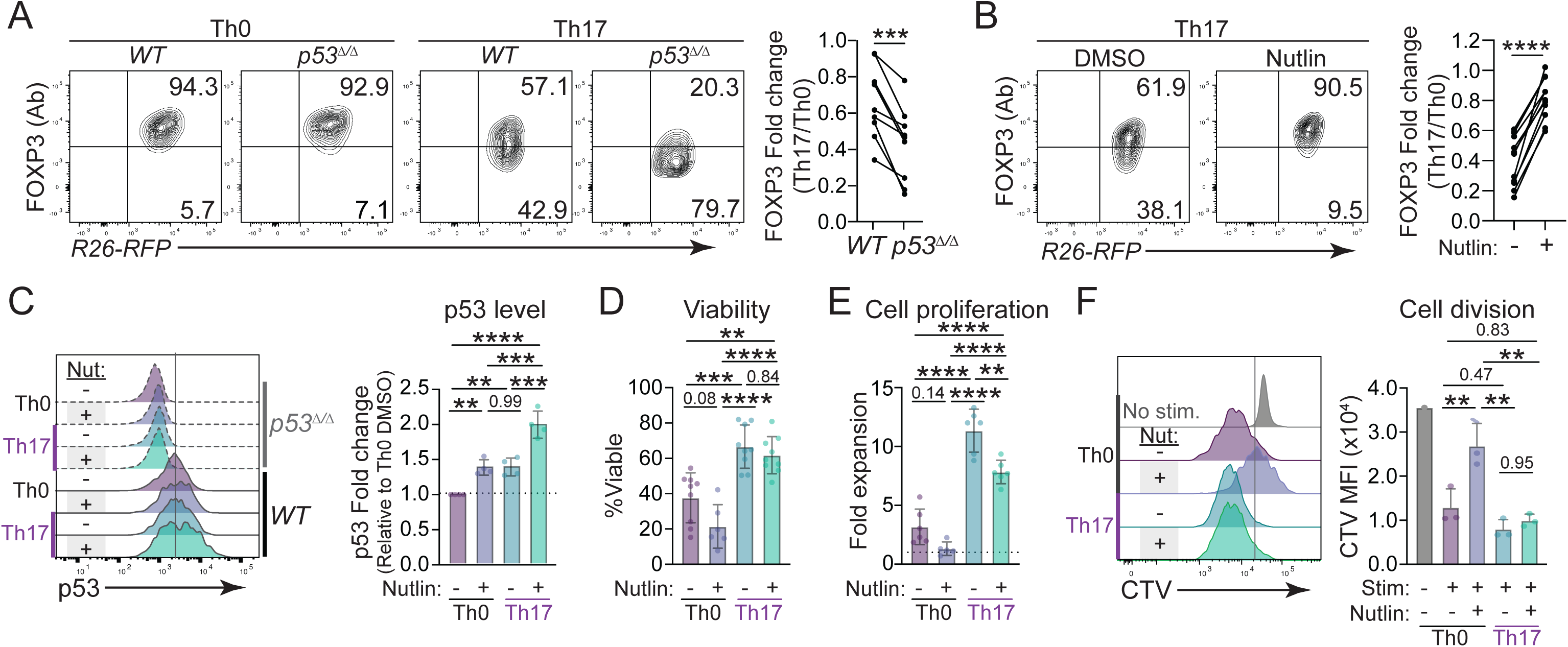
p53 activity promotes the maintenance of Foxp3 expression. (A) Representative flow plots (left) and paired quantification (right) of day 8 FOXP3 expression in WT and p53 Δ/Δ Tregs in Th17 and Th0 conditions. Data points represent biological replicates pooled from nine independent experiments. (B) Representative flow plots (left) and paired quantification (right) of day 8 FOXP3 expression in WT Tregs in Th17 conditions, treated with 10uM Nutlin or DMSO for 36 hours (starting day 3). Data points represent biological replicates pooled from ten independent experiments. (C) Representative histograms (left) and relative quantification (right) of p53 expression in Tregs in Th17 and Th0 conditions 24 hours after treatment with 10uM Nutlin or DMSO (starting day 3). Data points represent biological replicates pooled from four independent experiments. (D-E) Frequency of viable cells (D) and fold expansion of (E) Tregs in Th0 or Th17 conditions, treated with 10uM Nutlin or DMSO for 36 hours (starting day 3). Data points represent biological replicates pooled from four to six independent experiments. (F) Representative histogram (left) and MFI (right) of CellTrace proliferation dye for Tregs in Th0 or Th17 conditions, treated with 10uM Nutlin or DMSO for 36 hours (starting day 3). Tregs labeled with proliferation dye after washing out Nutlin on day 5, analysis on day 8. Data from n=3 mice per group. *p<0.05, **p<0.01, ***p<0.001, ****p<0.0001 by paired Student’s t-test (A-B) or Ordinary one-way ANOVA (C-F), mean ± s.d.

Next, we sought to determine whether enforced p53 stabilization could increase the maintenance of Foxp3 expression in Tregs in Th17 conditions. To promote p53 stabilization, we treated Tregs with Nutlin-3, a small molecule MDM2 inhibitor, to prevent p53 degradation during the first 36 hours of Th17 cytokine treatment. Strikingly, acute Nutlin treatment of *WT* Tregs fully preserved Foxp3 expression in Th17 conditions (Figure 3B). As expected, Nutlin treatment had no effect on improving Foxp3 maintenance in *p53^Δ/Δ^* Tregs, confirming that the activity of Nutlin required p53 protein stabilization (Extended Data Figure 3B). There was also no effect on *WT* Foxp3 levels in Th0 conditions (Extended Data Figure 3C). Interestingly, direct analysis of p53 protein levels by flow cytometry showed that treatment of Tregs with Th17 cytokines or Nutlin both increased p53 to similar levels (Figure 3C). However, Th17 cytokines and Nutlin together stabilized p53 protein levels to the greatest extent (Figure 3C). Therefore, p53 stabilization may protect Tregs from losing Foxp3 expression in Th17 conditions.

Since p53 is well-known for promoting apoptosis and cell cycle arrest, we investigated whether either of these mechanisms were responsible for the increased maintenance of Foxp3 in Tregs. For example, p53 activity could selectively prevent the proliferation and expansion of ex-Tregs, or it could cause the apoptosis of cells that lose Foxp3 expression. Both scenarios could lead to the selective maintenance of Foxp3-expressing cells over ex-Treg cells, increasing the apparent maintenance of Foxp3 expression with Nutlin treatment. However, there was no difference in the viability of Nutlin-treated versus DMSO (vehicle)-treated Tregs in Th17 conditions (Figure 3D).

While the fold expansion of Nutlin-treated Th17-conditioned Tregs was modestly, albeit significantly, reduced compared to vehicle-treated Th17-conditioned Tregs (Figures 3E), Tregs stained with a proliferation dye (CTV) after removing Nutlin from the media and then analyzed three days later, showed equivalent proliferation while maintaining Foxp3 expression (Figure 3F and Extended Data Figure 3D). Together, these results indicate that the selective apoptosis or cell cycle arrest of Tregs that lose Foxp3 are not the mechanisms by which p53 activity preserves Foxp3 expression in Tregs in Th17 conditions.

### p53 is not necessary for Foxp3 induction in T cells

To further understand the relationship between p53 and Foxp3 expression, we looked at the requirement for p53 during the development of natural Tregs (nTregs) in the thymus and induced Tregs (iTregs) *in vitro* (Figure 4A). In *Foxp3-GFP-hCre;p53^fl/fl^;R26^LSL-RFP^* (Treg.*p53^Δ/Δ^*) mice, frequencies of Tregs and Foxp3 expression levels in Tregs from the thymus were comparable to those in *WT* mice, indicating that p53 was not necessary to maintain Foxp3 expression immediately following Foxp3 induction after thymic development (Figures 4B-C). To determine whether p53 was necessary for TGF-β-induced iTreg generation *in vitro*, we isolated naïve CD4^+^ T cells from *WT* or *CD4-Cre;p53^fl/fl^* (T.*p53^Δ/Δ^*) mice, which delete p53 upon CD4 expression at the double positive stage of thymic development. In contrast to results from another group^40^, *p53^Δ/Δ^* iTregs developed normally, with comparable Foxp3 induction as with *WT* iTregs (Figures 4D-E). We confirmed p53-deficiency in cells by treatment with doxorubicin, a robust inducer of DNA damage and subsequent p53 stabilization, which showed strong induction of p53 protein in *WT* cells that was absent in *p53^Δ/Δ^*cells (Extended Data Figure 4A). We next tested whether stabilization of p53 through Nutlin treatment could increase Foxp3 induction in iTreg conditions. We found that Nutlin-treated T cells did not differentiate into iTregs more readily, nor did iTregs express higher levels of Foxp3 compared to DMSO-treated T cells, regardless of TGF-β treatment (Figures 4D-F). Therefore, p53 function does not appear to be necessary or sufficient for Foxp3 induction in T cells developing naturally in the thymus or induced by TGF-β treatment *in vitro*.

**Figure 4.**
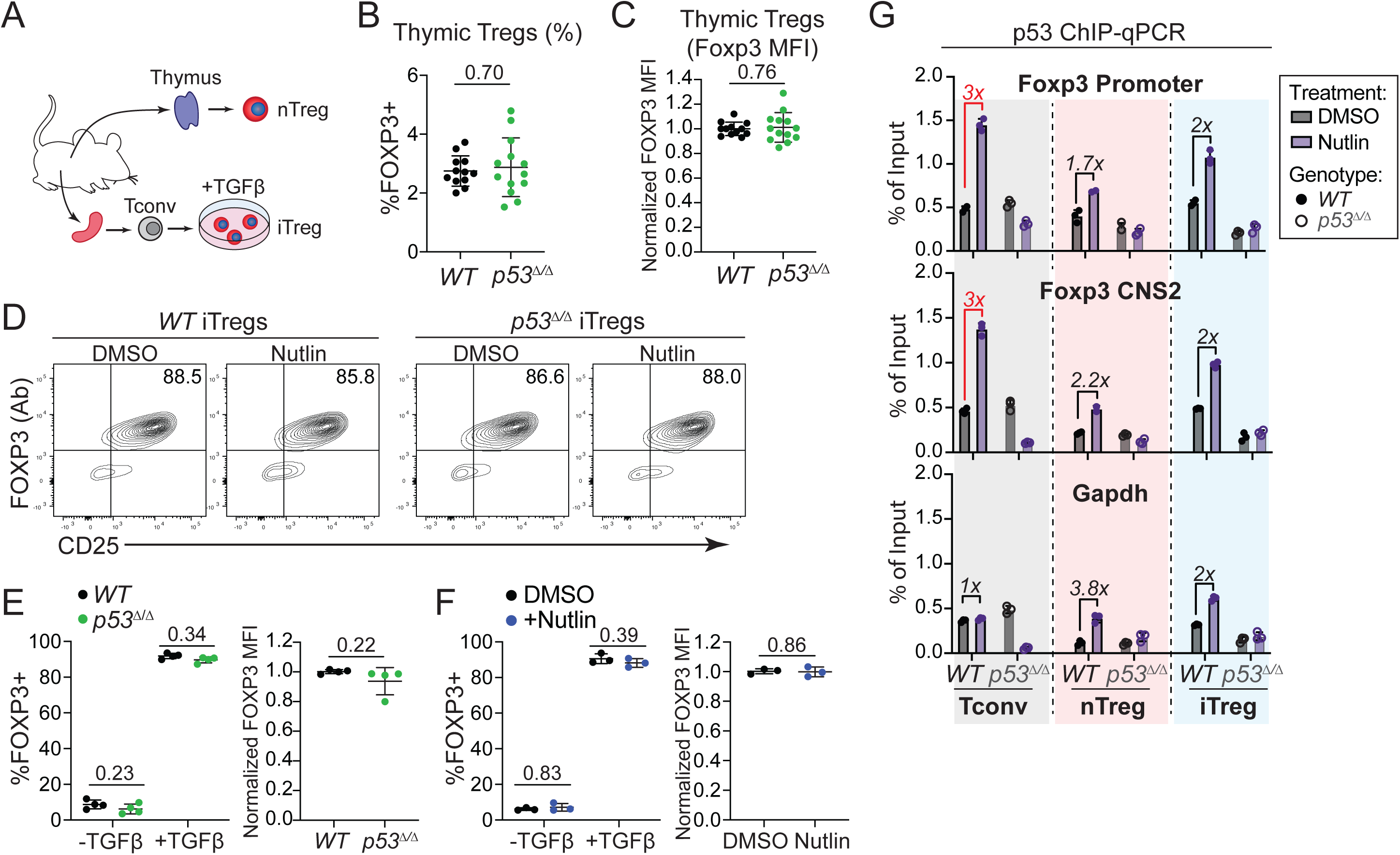
p53 is not necessary for Foxp3 induction in T cells. (A-B) Frequency (A) and FOXP3 MFI (B) of FOXP3+ Tregs cells in the thymus of WT and Treg.p53 Δ/Δ (p53 Δ/Δ) mice. Data from n=12-13 mice per genotype. (C) Representative flow plots and frequencies of iTreg generation from WT and p53 Δ/Δ naïve CD4+ T cells treated with 5 ng/mL TGF-b for 4 days and 10uM Nutlin or DMSO. (D) Frequency (left) and relative FOXP3 expression (right) of FOXP3+ iTregs from WT and p53 Δ/Δ naïve CD4+ T cells treated with 5 ng/mL TGF-b for 4 days. Data points represent biological replicates pooled from three independent experiments. (E) Frequency (left) and relative FOXP3 expression (right) of FOXP3+ iTregs from WT naïve CD4+ T cells treated with 5 ng/mL TGF-b for 4 days and 10uM Nutlin or DMSO. Data points represent biological replicates pooled from two independent experiments. (F) ChIP-qPCR analysis of indicated regions of Foxp3 locus or Gapdh in WT or p53 Δ/Δ conventional CD4+ T cells (Tconv), sorted Tregs (nTreg), or induced Tregs (iTreg), treated with 10uM Nutlin or DMSO for 36 hours. Numbers indicate fold change relative to WT DMSO condition. Data points represent technical replicates, representative of at least two independent experiments. For all plots, *p<0.05, **p<0.01, ***p<0.001, ****p<0.0001 by unpaired Student’s t-test, mean ± s.d.

Direct binding of p53 to the Foxp3 locus has also been reported in cancer cells as well as conventional CD4^+^ T cells, specifically at the promoter and CNS2 regions of the *Foxp3* locus^40,65^. To determine if p53 also binds to the *Foxp3* locus in Tregs, we performed p53 ChIP-qPCR on DMSO- or Nutlin-treated Tconv, nTregs, and iTregs from *WT* and T.*p53^Δ/Δ^* mice (Figure 4G). Again, Nutlin treatment boosted p53 protein levels in all *WT* cells but did not affect Foxp3 protein levels (Extended Data Figures 4B-C). Importantly, p53 stabilization in *WT* Tconv cells resulted in detectably increased p53 binding at both the *Foxp3* promoter (3x more) and *Foxp3* CNS2 (3x more) regions with Nutlin treatment, which was lost in *p53^Δ/Δ^* cells, as was previously reported (Figure 4G)^40^. However, despite increased p53 protein levels with Nutlin treatment in Tconv cells, nTregs and iTregs showed levels of p53 binding at the *Foxp3* promoter and *Foxp3* CNS2 region that did not exceed the background levels of p53 binding at the control *Gapdh* locus (Figure 4G). Thus, while these experiments confirmed prior findings that p53 binds the *Foxp3* locus in Tconv cells, we show here, and for the first time, that p53 does not appear to bind strongly to the *Foxp3* locus upon p53 stabilization in nTregs or iTregs. These data suggest that p53-mediated Foxp3 maintenance may not be the result of direct regulation of Foxp3 expression by the p53 transcription factor.

### Treg-specific p53 deficiency leads to increased ex-Treg frequencies *in vivo* in the Th17 cytokine-rich colon tissue

Since *p53^Δ/Δ^* Tregs were more prone to losing Foxp3 expression *in vitro*, we investigated whether *p53*-deficiency also affected the maintenance of Foxp3 expression *in vivo*. Using the Treg lineage tracing mice (Figure 1A), we assessed the frequency of Tregs (GFP^+^RFP^+^) and ex-Tregs (GFP^-^RFP^+^) in lymphoid and non-lymphoid tissues from *WT* and Treg.*p53^Δ/Δ^* mice. There were no significant differences in Treg frequencies between *WT* and Treg.*p53^Δ/Δ^*mice in any of the tissues analyzed (Figures 5A-B and Extended Data Figure 5A). While ex-Treg frequencies were similar in the lymph nodes, spleen, and lungs in Treg.*p53^Δ/Δ^* mice, ex-Treg frequencies in the colon were significantly increased (Figures 5A, 5C-D and Extended Data Figures 5B-C). Interestingly, analysis of Neuropilin 1 (Nrp1) and ROR-γt expression in the ex-Tregs from the colon showed that the ex-Tregs were significantly enriched only in the Nrp1^+^ cells (Extended Data Figures 5D-E). This may suggest that the increased ex-Tregs observed in the Treg.*p53^Δ/Δ^* mice are derived from Nrp1^+^ nTregs and not ROR-γt^+^ iTregs. Collectively, these results show that Treg-specific *p53*-deficiency results in increased ex-Treg frequencies specifically in the colon, which is likely due to an impaired maintenance of nTregs in tissues with Th17 associated cytokines.

**Figure 5.**
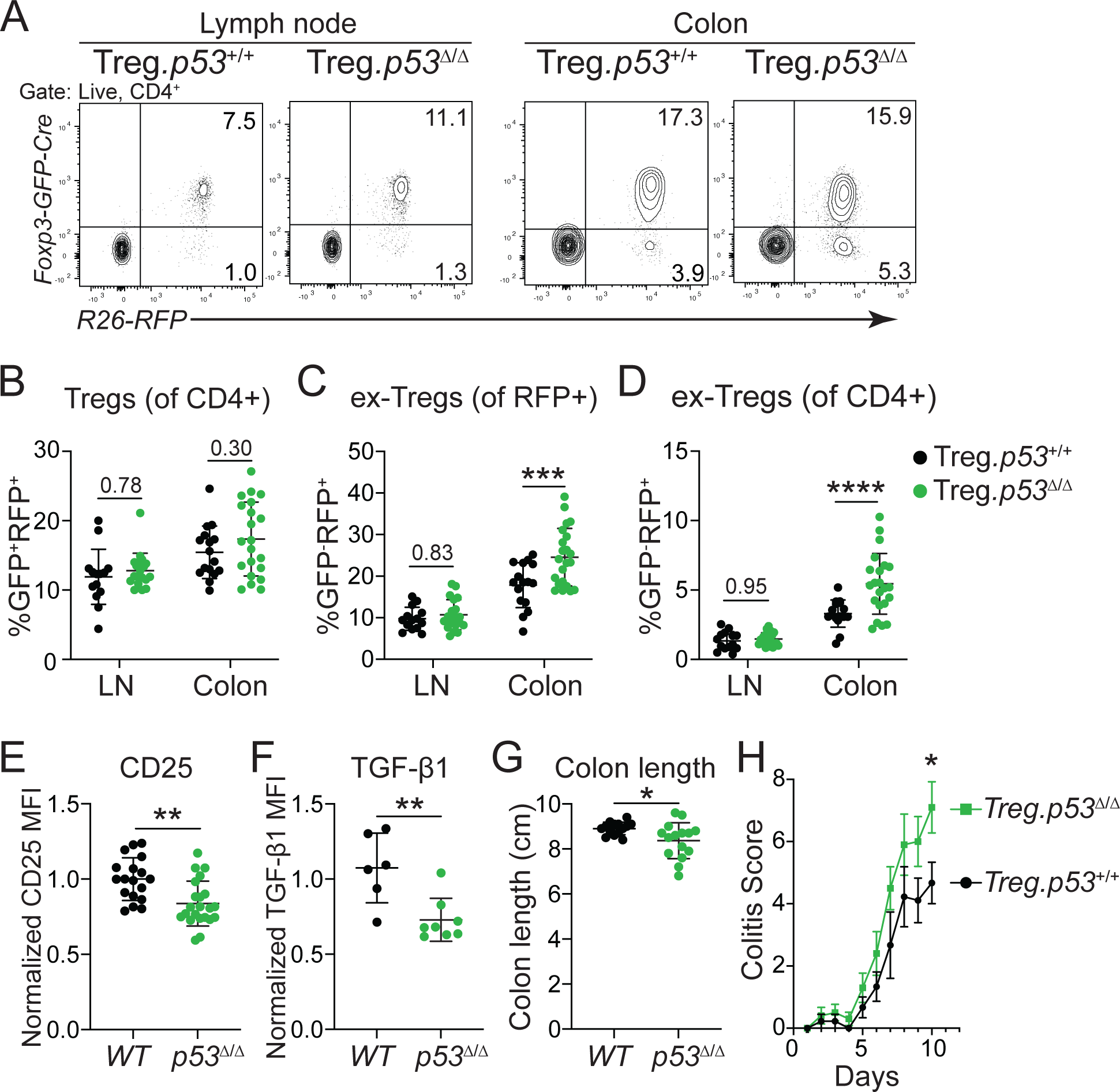
Treg-specific p53 deficiency leads to increased ex-Treg frequencies in vivo in Th17 cytokine-rich colon tissues. (A) Representative flow plots and frequencies of Tregs and ex-Tregs in the lymph node (LN) and colon of WT and Treg.p53 *Δ/Δ* mice. (B) Frequency of GFP+RFP+ Tregs in the LN and colon of WT and Treg.p53 Δ/Δ mice. Data from n=14-21 mice per genotype. (C-D) Frequency of ex-Tregs of RFP+ (C) or CD4+ (D) cells in the LN and colon of WT and Treg.p53 Δ/Δ mice. Data from n=14-21 mice per genotype. (E) Normalized CD25 MFI of GFP+RFP+ Tregs from WT and Treg.p53 Δ/Δ (p53 Δ/Δ) mice. Data from n=18-21 mice per genotype. (F) Normalized TGF-b1/LAP MFI of GFP+RFP+ Tregs from WT and Treg.p53 Δ/Δ (p53 Δ/Δ) mice. Data from n=6-8 mice per genotype. (G) Measurements of colons from WT and Treg.p53 *Δ/Δ* (p53 Δ/Δ) mice. Data from 15-16 mice per genotype. (H) Combined DSS colitis scores over time for WT versus Treg.p53 Δ/Δ mice. Data from n=9-10 mice per genotype, representative of two experiments. *p<0.05, **p<0.01, ***p<0.001, ****p<0.0001 by two-way ANOVA (B-D,H), or unpaired Student’s t-test (E-G), mean ± s.d.

Phenotypic analysis of the Tregs in the colon revealed that *p53^Δ/Δ^*Tregs had decreased expression of key functional surface markers, such as CD25 and TGF-β1 (Figures 5E and 5F). Furthermore, the colons of Treg.*p53^Δ/Δ^* compared to *WT* mice were significantly shorter, suggestive of increased intestinal inflammation in these mice (Figure 5G). Finally, we tested the consequences of Treg-specific *p53*-deficiency in the control of an induced model of intestinal inflammation using low-dose dextran sulfate sodium (DSS) in the drinking water to induce colitis. Treg.*p53^Δ/Δ^* mice had a more severe colitis compared to *WT* mice (Figure 5H). Therefore, mice with *p53*-deficient Tregs exhibit functional Treg impairment and increased ex-Treg generation in the colon, leading to diminished control of intestinal inflammation.

## Discussion

Many studies have illustrated the importance of epigenetic modifications in supporting Treg function^4,7–18,66–70^. The repressive chromatin modification H3K27me3 has been shown to act as a switch for Treg stability, but the pathways regulated by H3K27me3 in Tregs have yet to be uncovered^16,19–24,71^. In this study, we found an enrichment for genes regulated by *Trp53*, or p53, among differentially expressed genes between *Ezh2^Δ/Δ^* or *Jmjd3^Δ/Δ^* Tregs versus *WT* Tregs. P53 target genes were largely upregulated in *Ezh2^Δ/Δ^* Tregs and downregulated in *Jmjd3^Δ/Δ^*Tregs, demonstrating that the loss or gain of epigenetic repression by H3K27me3 modifications may underlie p53 regulation. Among the most differentially expressed genes was *Cdkn2a*, whose protein product p19ARF negatively regulates MDM2, an E3 ubiquitin ligase responsible for maintaining low levels of p53 protein in healthy cells. Other studies have shown that loss of *Ezh2* and H3K27me3, either through genetic deficiency or pharmacological inhibition of EZH2, led to increased p53 accumulation due to de-repression of *Cdkn2a*/p19ARF in various cell types^72–74^. Thus, while it might be expected that elevated p19ARF protein levels in *Ezh2^Δ/Δ^*Tregs would promote increased p53 activity, our finding that p53 is activated in this way in a non-cancerous cell type under potentially stressful environments is unique. We hypothesize that the *Cdkn2a*/p19ARF locus may more generally act as a sensor for cells that are losing their differentiated identity due to a global loss of H3K27me3 modifications. Therefore, triggering p53 activation may serve to stabilize cells that would otherwise lose their cellular identity. It is possible that this genetic circuit evolved for this purpose in all cell types, even though it is recognized prominently for its role in tumor suppression by cell cycle arrest or apoptosis of transformed cancer cells.

Nevertheless, p53 has also been described as a guardian against autoimmunity, as mice deficient in all cells for p53 are not only more susceptible to cancer, but also several autoimmune diseases^36–40^. However, exactly which cell types contribute to the exacerbated autoimmunity seen in global and T cell-specific p53-deficient mice is unclear. Studies that have implicated Tregs have focused on their impaired differentiation from *p53^Δ/Δ^* CD4^+^ T cells as the source of increased susceptibility to autoimmunity^40,41^. Unexpectedly, we found no defect with *in vitro* iTreg differentiation from *p53^Δ/Δ^* T cells. We sought to investigate the effect of p53 deficiency on the maintenance of Foxp3 and Treg identity unrelated to Treg differentiation by using mice whose cells lose p53 only after Foxp3 expression is induced. Our data suggests that p53 does not play a general role in Foxp3 maintenance, but that it is important in the context of Th17-polarizing cytokines, i.e. IL-1β, IL-6, and IL-23, which are often produced in gut-associated tissue environments. Other studies have reported that *p53* deficiency in T cells increases Th17 differentiation at the expense of Treg induction, supporting an antagonistic relationship between p53 and Th17 cytokines in regards to *Foxp3* expression^40,41^. Despite those findings, to our knowledge, p53 has not been previously identified as an important factor for Tregs in Th17 cytokine-rich tissues, such as the colon. In this study, we show that *p53^Δ/Δ^* colonic Tregs were more prone to losing *Foxp3* expression compared to their *WT* counterparts, and that those Tregs that retained *Foxp3* expression had lower levels of CD25 and TGF-β, two important mechanisms used by Tregs for their suppressive capacity^75,76^. Therefore, p53 deficiency in colonic Tregs likely compromises their function both by rendering them more vulnerable to cytokine-mediated reprogramming and impairing key mechanisms of suppression.

Most intriguingly, our study provides a new example of p53’s role - to restrict cell plasticity in a non-transformed or pluripotent cell type. As a tumor suppressor, p53 plays a critical role in restraining cell transformation into self-renewing cancer cells^77–80^. It has also been shown to inhibit somatic cell reprogramming into induced pluripotent stem cells (iPSCs)^81–84^. In both cases, p53 limits the de-differentiation of cells during transformation by strong oncogenes. Here, we show that stress caused by Th17 cytokines can also trigger p53 activation in Tregs, and that acute stabilization of p53 protein is sufficient to protect Tregs from losing Foxp3 expression and their immunosuppressive function. Thus, changes in cues received by cells across changing or different tissue environments can trigger p53 to support the maintenance of cellular identity. Furthermore, while p53 has long been known to operate as a tumor suppressor by activating apoptosis or cell cycle arrest in transformed cells, in this study we demonstrate that p53 can protect against Th17 cytokine-mediated FOXP3 loss independently of these two mechanisms *in vitro*^85–90^. We hypothesize that p53 may function more broadly as a mechanism for many differentiated cell types to preserve cell lineage identities, and that sensing H3K27me3 modifications at the *Cdkn2a* locus serves as a means to monitor cell identity broadly in many cell types and appropriately trigger p53 function when cells do not properly maintain this crucial repressive chromatin modification. Still, much work is required to understand how p53 activity can maintain Treg identity independently of direct *Foxp3* genetic regulation, or canonical pruning of destabilized cells by apoptosis or cell cycle arrest.

Regardless of this deeper mechanistic understanding of how p53 functions to maintain Treg function, this study reveals the potential for p53 stabilization in Tregs as a new therapeutic approach for treating intestinal inflammation or autoimmunity more generally. Broadly targeting p53 in autoimmune disease has been previously proposed due to its protective role in immune regulation, and other groups have shown that treatment with different small molecule activators of p53 can improve autoimmunity and boost Treg frequencies *in vivo*^41,91,92^. However, because of the possibility of promoting apoptosis or cell cycle arrest in cell types that are more sensitive to p53 activation, it may be more feasible to use p53-activating treatments in the context of adoptive Treg transfer or chimeric antigen receptor (CAR) Tregs^93–98^. In particular, Treg therapies for IBD may benefit from p53 activation *ex vivo* due to enhanced maintenance of function and stability of transferred Tregs that migrate into the gut-tissues post transfer. Further research will be necessary to directly address the potential benefit of p53 stabilization on Treg-targeted therapies.

In summary, our results show that p53 expression can be triggered by loss of H3K27me3 in Tregs and that p53 serves as a crucial protector of Treg identity in Th17-polarizing environments like the colon. These findings illustrate how Tregs can utilize epigenetic sensors to trigger mechanisms that can preserve Treg functions when encountering inflammatory cytokines upon entering diverse tissue environments. The identification of p53 as a key promoter of Treg function in colonic tissues may inform improved methods to prepare Treg-based therapies for advancing the treatment of inflammatory bowel diseases.

## Materials and Methods

### Mice

All mice used were bred onto a C57BL/6 background for a minimum of ten generations. All mouse and *in vitro* experiments used comparisons between littermates or age-matched control mice (12–20 weeks old) and included mice of both sexes. *Foxp3-GFP-hCre* mice were kindly provided by Dr. Bluestone (University of California, San Francisco; JAX:023161) and the *R26^LSL-RFP^* allele was described previously^99^. *CD4-Cre* mice were obtained from Jackson laboratories (JAX:022071) and bred in house. *Ezh2^fl^* mice harbor loxP sites flanking exons 16–19 encoding the SET domain^100^. The *p53*^fl^ allele originated from *Foxp3^GFP-DTR^;Kras*^G12D^*;p53*^fl/fl^ mice that were kindly provided by Dr. Jacks (Massachusetts Institute of Technology) and was crossed to *Foxp3-GFP-hCre or CD4-Cre mice to generate Foxp3-GFP-hCre;p53^fl/fl^ and CD4-Cre;p53^fl/fl^ mice*.

### Cell isolation for flow cytometry

Single cell suspensions from lymphoid organs were prepared by mechanical disruption in an ice-cold PBS buffer containing 2% FBS and 4 mM EDTA (Invitrogen) and passing through 40 μm filters. Spleens were subsequently subjected to red blood cell lysis using ACK buffer (150mM NH4Cl, 10mM KHCO3, 0.1mM Na2EDTA, pH7.3). For isolation of lymphocytes from the colon, the large intestine was first resected by cutting below the cecum and above the anus. After removal of fat, the colon was flushed with ice-cold PBS to remove intestinal contents before being cut longitudinally and rinsed with PBS. Intestine was cut into 1-2 cm pieces and placed with a stir bar in 50 mL flasks with pre-warmed Hank’s balanced salt solution (HBSS) containing 10 mM HEPES (Thermofisher Scientific), 2 mM L-glutamine, 1 mM dithiothreitol (DTT; Fisher Scientific), and 1.25 mM EDTA and incubated for 20 minutes on a stir plate at 37’C. Tissues were then strained into 100 μm filters, rinsed with PBS, transferred to conical tubes containing PBS and then shaken vigorously and strained back into 100 μm filters. Pieces were returned to their original flasks with pre-warmed RPMI containing 25 mM HEPES, 2 mM L-glutamine, 1 mg/mL Collagenase VIII (Sigma-Aldrich), and 5 μg/mL DNase I (Roche) and incubated for 30-45 minutes on a stir plate at 37’C. After digestion, samples were filtered through 100 μm filters into 50 mL conical tubes, with remaining pieces of tissue broken up using the plunger of a syringe and RPMI wash buffer containing 5%FCS, 25 mM HEPES, and 2 mM L-glutamine added to stop the reaction. After centrifugation, cells were resuspended in 44% Percoll (Cytiva), transferred to FBS-coated 15 mL conical tubes, underlaid with 67% Percoll, and centrifuged at 2800 rpm for 20 minutes at room temperature. Lymphocytes were collected at the Percoll interphase and washed with RPMI wash buffer before staining. For isolation of lymphocytes from lungs, RPMI digestion media containing 25 mM HEPES, 20 μg/mL DNase I, and 16 mg/mL Collagenase D (Roche) was injected into the trachea to inflate the lungs before resection. Lungs were then chopped and incubated in digestion media for 45 minutes at 37’C with gentle shaking. Digested tissue was then filtered through 70 μm filters into 50 mL conical tubes, with remaining pieces of tissue broken up using the plunger of a syringe. After centrifugation, cells were subjected to red blood cell lysis using ACK buffer and washed with HBSS before staining.

### Flow cytometry

Flow cytometry was performed on a BD LSR Fortessa X20 (BD Biosciences) and datasets were analyzed using FlowJo software (Tree Star). Dead cells were first stained with Live/Dead Fixable Blue or Aqua Dead Cell Stain kit (Molecular Probes) in PBS for 15 min at 4^◦^C. Cell surface stains were carried out for 30 min at 4^◦^C using a mixture of fluorophore-conjugated antibodies: anti-mouse CD45 (30-F11, BioLegend), anti-mouse CD4 (RM4-5, BioLegend), anti-mouse CD8a (53–6.7, BioLegend), anti-mouse CD25 (PC61, BioLegend), anti-mouse Nrp1 (3DS304M, Thermo Scientific) and anti-mouse LAP/TGF-β1 (TW7-16B4, BioLegend). Cells were fixed in 4% paraformaldehyde for 15 minutes at room temperature followed by permeabilization with 0.1% Triton X-100 for 20 minutes at room temperature. Intracellular staining was performed in PBS containing 0.5% BSA using anti-mouse Foxp3 (FJK-16S, Thermo Scientific), anti-mouse RORγt (Q31-378, BD Biosciences), and anti-mouse p53 (1C12, Cell Signaling Technology) for 1 hour at 4’C.

### *In vitro* Treg reprogramming

Single cell suspensions were generated from mice and enriched for CD4+ T cells by negative selection using EasySep magnetic bead kit (STEMCELL Technologies). Enriched cells were stained with anti-mouse CD4 (RM4-5, BioLegend), anti-CD25 (PC61, BioLegend) and anti-CD62L (MEL1-14, Biolegend) in Opti-MEM containing 2% FBS. Naïve Treg cells (CD4^+^CD25^+^Foxp3^GFP+^R26^RFP+^CD62L^+^) were sorted using an Aria Fusion sorter (BD Biosciences) with a 70μm nozzle. Sorted Tregs were cultured in DMEM medium supplemented with 10% FBS (BioWest), 1% non-essential amino acids, 1 mM sodium pyruvate, 2 mM L-glutamine, 10 μM HEPES and 55 μM β-mercaptoethanol with 2000 U/mL IL-2 and CD3/CD28 Dynabeads (Gibco) for 2.5 days. Cells were then treated with 200 U/mL IL-2 alone or in combination with either 20 ng/mL IL-12, 20 ng/mL IL-4, or 20 ng/mL IL-6, 20 ng/mL IL-23 and 10 ng/mL IL-1β for 5 days, with more cytokines added after 2 days. In experiments involving Nutlin, cells received 10 uM Nutlin-3 (VWR) or DMSO for the first 36 hours that pro-inflammatory cytokines were present, with cytokine-containing medium being replaced for all samples upon wash out. For cell proliferation experiments, Tregs were labeled with CellTrace Far Red Cell Proliferation Kit (Thermo Fisher Scientific) after Nutlin wash out and analyzed for dilution of the dye 3 days later.

### Western blotting

Cells were collected and lysed in Laemmli buffer (31.5 mM Tris-HCl, pH 6.8, 10% glycerol, 1% SDS, 0.005% Bromophenol Blue) before being boiled for 10 minutes. Samples were then run on a 4-20% Mini-Protean TGX gel (Bio-Rad) and transferred to PVDF membranes. Membranes were incubated in Intercept PBS blocking buffer (Li-Cor) for 1 hour at room temperature with shaking before primary staining with anti-p53 (1C12, Cell Signaling Technology), anti-p19ARF (5-C3-1, Thermo Scientific), or anti-β-actin (13E5, Cell Signaling Technology) overnight at 4’C with shaking. After three 10-minute washes with TBST buffer (Tris-buffered saline, 0.1% Tween 20), membranes were stained with anti-mouse, anti-rat, or anti-rabbit secondary antibodies (Li-Cor) for 1 hour at room temperature with shaking and imaged using an Odyssey CLx imager (Li-Cor).

### RT-qPCR

Naïve Tregs (CD4^+^CD25^+^Foxp3^GFP+^R26^RFP+^CD62L^+^) were sorted from *Foxp3-GFP-hCre;R26^RFP^*, *Foxp3-GFP-hCre;Ezh2^fl/fl^;R26^RFP^*, and *Foxp3-GFP-hCre;Ezh2^fl/fl^;p53^fl/fl^;R26^RFP^*mice and cultured for 4 days in DMEM medium supplemented with 10% FBS (BioWest), 1% non-essential amino acids, 1 mM sodium pyruvate, 2 mM L-glutamine, 10 μM HEPES and 55 μM β-mercaptoethanol with 2000 U/mL IL-2 and CD3/CD28 Dynabeads. Upon collection, total RNA was isolated using the RNeasy Mini kit (Qiagen). cDNA was subsequently synthesized using the iScript cDNA Synthesis kit (Bio-Rad). RT-qPCR was performed using Fast SYBR Green Master Mix (Applied Biosystems) and a QuantStudio 6 Pro Real-Time PCR machine (Applied Biosystems). Relative gene expression was determined by the 2^-Δ^ ^ΔCt^ method.

### Chromatin Immunoprecipitation-qPCR

Naïve Tregs (CD4^+^CD25^+^CD62L^+^GITR^+^) and Tconv (CD4^+^CD25^-^CD62L^+^GITR^-^) were first sorted from *CD4-Cre;p53^+/+^* or *CD4-Cre;p53^fl/fl^* mice before being cultured on plates pre-coated with anti-CD3 and anti-CD28 in DMEM medium supplemented with 10% FBS (BioWest), 1% non-essential amino acids, 1 mM sodium pyruvate, 2 mM L-glutamine, 10 μM HEPES and 55 μM β-mercaptoethanol with 200 U/mL IL-2. Some naïve Tconv were also cultured with 5 ng/mL TGF-β to generate induced Tregs (iTreg). Cells were collected after 4 days, with all populations having received either DMSO or 10 uM Nutlin-3 for the last 36 hours of culture. Upon collection, cells were fixed and crosslinked with 2 mM disuccinimidyl glutarate (DSG) and 0.9% formaldehyde for p53 ChIP experiments. Chromatin immunoprecipitation was performed with anti-p53 (1C12, Cell Signaling) bound to Protein A/G magnetic beads (Pierce) using the iDeal ChIP-qPCR kit (Diagenode) according to the manufacturer’s instructions. Samples were sheared using a Covaris S2 bioruptor. RT-qPCR was performed using Fast SYBR Green Master Mix (Applied Biosystems) and a QuantStudio 6 Pro Real-Time PCR machine (Applied Biosystems). %Input was calculated using the formula: %Input=2^[(Ct_input_-log2(X))-Ct_sample_]*100%, where X is the input dilution.

For H3K27me3 ChIP experiments, Tregs (CD4^+^CD25^+^Foxp3^GFP+^R26^RFP+^CD62L^+^) from *Foxp3-GFP-hCre;R26^RFP^*, *Foxp3-GFP-hCre;Ezh2^fl/fl^;R26^RFP^*, and *Foxp3-GFP-hCre;Jmjd3^fl/fl^;R26^RFP^*mice were sorted and cultured for 4 days in DMEM medium supplemented with 10% FBS (BioWest), 1% non-essential amino acids, 1 mM sodium pyruvate, 2 mM L-glutamine, 10 μM HEPES and 55 μM β-mercaptoethanol with 2000 U/mL IL-2 and CD3/CD28 Dynabeads (Gibco). Upon collection, cells were fixed and crosslinked with 0.9% formaldehyde. Chromatin immunoprecipitation was performed with anti-H3K27me3 (C36B11, Cell Signaling Technology) or control IgG antibody (2729, Cell Signaling Technology) using the same materials and equipment as above.

### DSS Colitis

Dextran sulfate sodium (DSS) salt was dissolved to a final concentration of 1.5% w/v in autoclaved drinking water and provided to experimental mice for 10 days, with fresh DSS-supplemented water replacement every two days. Mice were scored daily based on stool consistency, presence of blood in stool, and weight loss, using the following scoring system:

Stool consistency scoring: 0 – Normal stool consistency, 2 – Soft stools (no adhesion to the anus), 4 – Diarrhea (adhesion to the anus)

Blood in stool scoring: 0 – No blood in stool, 2 – Traces of blood in stool/Positive hemoccult, 4 – Gross rectal bleeding

Weight loss scoring: 0 – No weight loss, 1 – 1-5% loss, 2 – 6-10% loss, 3 – 11-15% loss, 4 – >15% loss

### CUT&Flow

Naïve Tregs (CD4^+^CD25^+^Foxp3^GFP+^R26^RFP+^CD62L^+^) from *Foxp3-GFP-hCre;R26^RFP^* mice were sorted and cultured for 4 days in DMEM medium supplemented with 10% FBS (BioWest), 1% non-essential amino acids, 1 mM sodium pyruvate, 2 mM L-glutamine, 10 μM HEPES and 55 μM β-mercaptoethanol with 2000 U/mL IL-2 and CD3/CD28 Dynabeads (Gibco). Upon collection, cells were lysed and nuclei extracted in nuclear extraction buffer (20 mM <u>HEPES</u> pH 7.9, 10 mM KCl, 0.1% Triton X-100, 20% glycerol, 0.5 mM spermidine and 1x Roche, cOmplete, Mini, EDTA-free Protease Inhibitor Cocktail) for 10 min on ice. Each reaction was incubated with primary anti-H3K27me3 antibody (C36B11, Cell Signaling Technology) in antibody 150 buffer (2 mM EDTA in Digitonin 150 buffer) for 1h at RT, followed by incubation with secondary antibody (Epicypher) in Digitonin 150 buffer (20 mM HEPES pH 7.5, 150 mM NaCl, 0.5 mM spermidine, 1x protease inhibitor cocktail and 0.01% digitonin) for 30 min at RT. Nuclei were next washed once in Digitonin 150 buffer and incubated with pAG-Tn5 (Epicypher) in Digitonin 300 buffer (20 mM HEPES pH 7.5, 300 mM NaCl, 0.5 mM spermidine, 1x protease inhibitor cocktail and 0.01% digitonin) for 1h at RT. After one wash in Digitonin 300 buffer, pAG-Tn5 was activated and tagmentation performed by incubation in tag-mentation buffer (20 mM HEPES pH 7.5, 300 mM NaCl, 0.5 mM spermidine, 1x protease inhibitor cocktail and 10 mM MgCl_2_) for for 1h at 37^◦^C. Tagmented nuclei were combined back into one tube per sample and incubated in TAPS buffer (10 mM TAPS pH 8.5 and 0.2 mM EDTA) for 5 min at RT to stop the tagmentation reaction. Samples were then resuspended in Krishan solution (3.8 mM sodium citrate; 46 µg/mL propidium iodide (PI); 0.01% NP-40; 10 µg/mL RNase A) and kept at 4^◦^C, protected from light, for up to three days, until sorting.

PI-stained nuclei were sorted based on DNA content in an Astrios EQ instrument (Beckman Coulter). Five fractions were defined: 0-G1, G2-M and three equal subdivisions of the S phase, classified as Early S, Mid S and Late S. After sorting, samples were incubated in SDS release buffer (10 mM TAPS pH 8.5 and 0.1% SDS) for nuclei lysis and release of tagmented chromatin for 1h at 58^◦^C. SDS was neutralized by addition of SDS quench buffer (0.67% Triton X-100 in nuclease-free H2O) and PCR performed with NEBNext High-Fidelity 2x PCR Master Mix (NEB, M0541L) and barcoded primers for library construction, with the following protocol: 5 min at 58^◦^C, 5 min at 72^◦^C, 45 s at 98^◦^C, 18 cycles of 15 s at 98^◦^C and 10 s at 60^◦^C, followed by 1 min at 72^◦^C for final extension.

### Statistical Methods

p values were obtained from unpaired two-tailed Student’s t tests for all statistical comparisons between two groups, and data were displayed as mean土 SD. For multiple comparisons, one-way ANOVA or two-way ANOVA was used. For DSS colitis disease progression, two-way ANOVA was used with Sidak’s multiple comparisons test performed at each time point or by multiple regression analysis. p values are denoted in figures by *p < 0.05, **p < 0.01, ***p < 0.001, and ****p < 0.0001.

## Supporting information

Silveria S et al. Supplement

## Acknowledgments

We thank Hector Nolla, Alma Valleros, Kartoosh Heydari and Melaine Delcroix of the UC Berkeley Cancer Research Laboratory Flow Cytometry Facility. We also thank all members of the DuPage lab for providing feedback on the research approach and critically reviewing the manuscript. This research was supported by the National Institutes of Health (1DP2CA247830-01 to M.D. and R35GM156411 to S.R.), an IVRI-Aduro Biotech sponsored research award (M.D.), and the Siebel Stem Cell Institute (M.D.), an American Cancer Society grant RSG-22-026-01 (S.R.), and the RNA Bioscience Initiative (University of Colorado School of Medicine). S.S. was supported by a Cancer Research Coordinating committee (CRCC) fellowship. S.R. is a Pew-Stewart Scholar. M.D. is a Pew-Stewart Scholar and a St. Baldrick’s Scholar with generous support from Hope with Hazel.

