## Supplementary material for "Epigenetically regulated p53 activity maintains intestinal regulatory T cell identity to prevent inflammation": Silveria S et al. Supplement

### Supp Figure 1: Regulation of H3K27me3 controls Foxp3 maintenance in Tregs

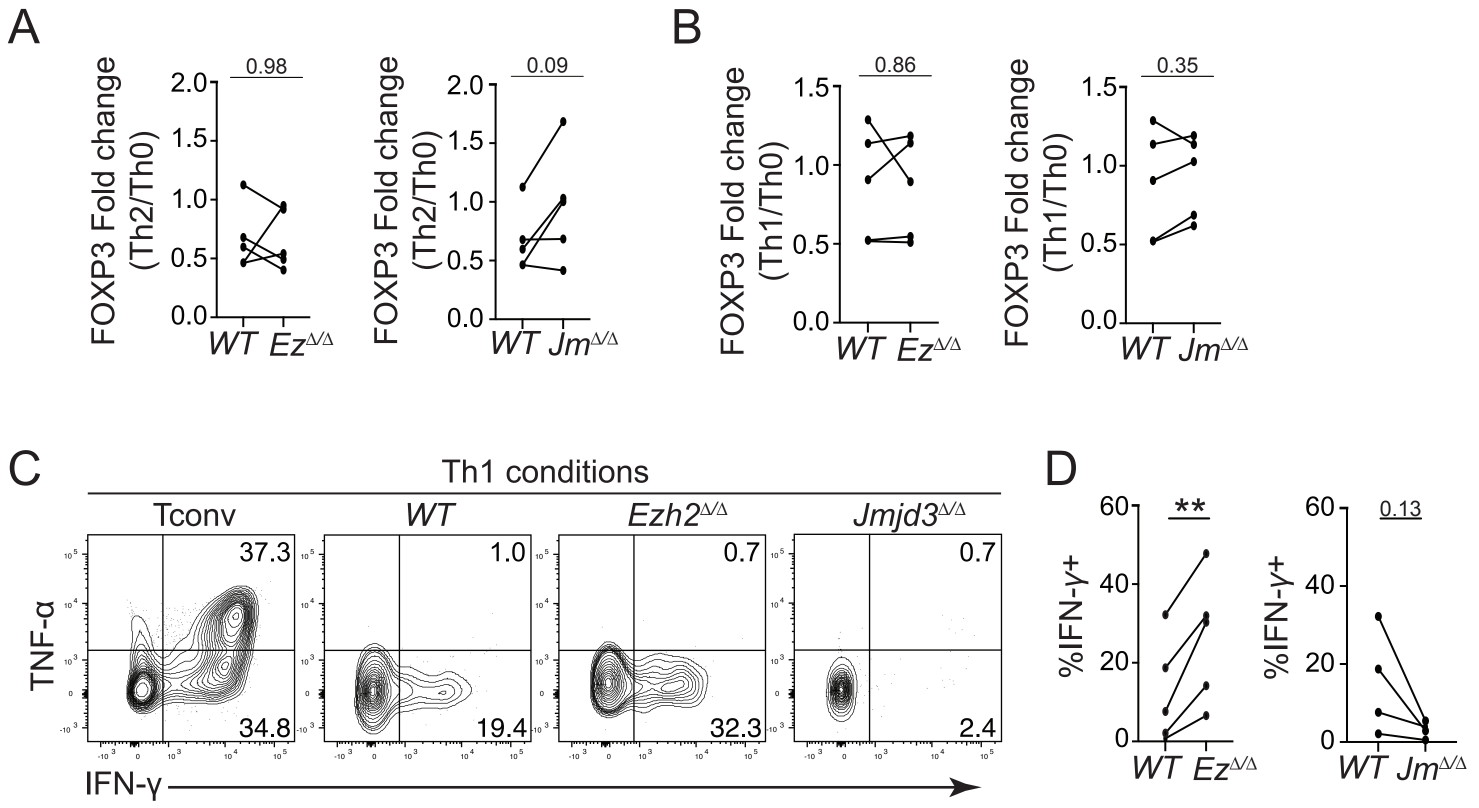


Supp. Figure 1 | Regulation of H3K27me3 controls Foxp3 maintenance in Tregs. (A) Paired quantification of FOXP3 fold change in Th2 relative to Th0 conditions for WT versus Ez Δ/Δ (left) or Jm Δ/Δ (right) Tregs. Data points represent biological replicates pooled from four independent experiments. (B) Paired quantification of FOXP3 fold change in Th1 relative to Th0 conditions for WT versus Ez Δ/Δ (left) or Jm Δ/Δ (right) Tregs. Data points represent biological replicates pooled from four independent experiments. (C) Representative flow plots for IFN-γ expression in WT, Ezh2 Δ/Δ, or Jmjd3 Δ/Δ Tregs and Tconv in Th1 conditions. (D) Paired frequency of IFN-γ+ cells in Th1 conditions for WT versus Ez Δ/Δ (left) or Jm Δ/Δ (right) Tregs. Data points represent biological replicates pooled from four to five independent experiments. For all plots, *$p<0.05$, **$p<0.01$, ***$p<0.001$, ****$p<0.0001$ by paired Student's t-test, mean ± s.d.

Supp Figure 2: Altered H3K27me3 levels control the activation of the p53 pathway in Tregs

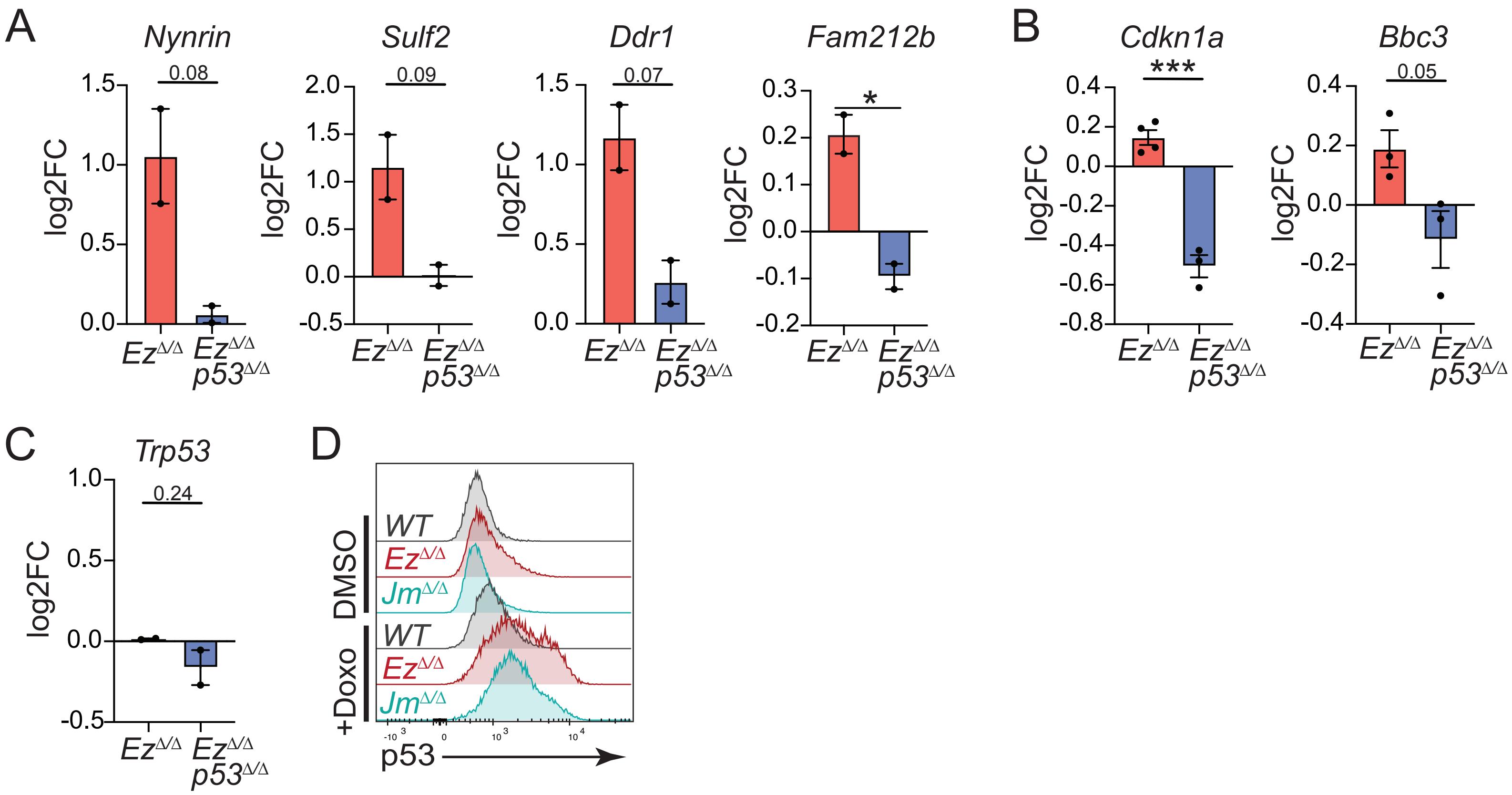


Supp. Figure 2 | Altered H3K27me3 levels control the activation of the p53 pathway in Tregs. (A) qPCR log2 fold change analysis of select p53 targets in Ezh2 Δ/Δ (EzΔ/Δ) or Ezh2 Δ/Δp53Δ/Δ (EzΔ/Δp53Δ/Δ) versus WT Tregs activated 4 days in vitro. Data points represent biological replicates pooled from two to three independent experiments. (B) qPCR log2 fold change analysis of canonical p53 targets in Ezh2 Δ/Δ (EzΔ/Δ) or Ezh2 Δ/Δp53Δ/Δ (EzΔ/Δp53Δ/Δ) versus WT Tregs activated 4 days in vitro. Data points represent biological replicates pooled from two to three independent experiments. (C) Representative histograms of p53 expression for 4 day in vitro-activated WT, EzΔ/Δ, and Jm Δ/Δ Tregs treated with DMSO or 50 nM doxorubicin (doxo) for 4 hours. (D) qPCR log2 fold change analysis of Trp53 in EzΔ/Δ or EzΔ/Δp53Δ/Δ versus WT Tregs activated 4 days in vitro. For all plots, *p<0.05, **p<0.01, ***p<0.001, ****p<0.0001 by unpaired Student’s t-test, mean ± s.d.

Supp Figure 3: p53 activity promotes the maintenance of Foxp3 expression

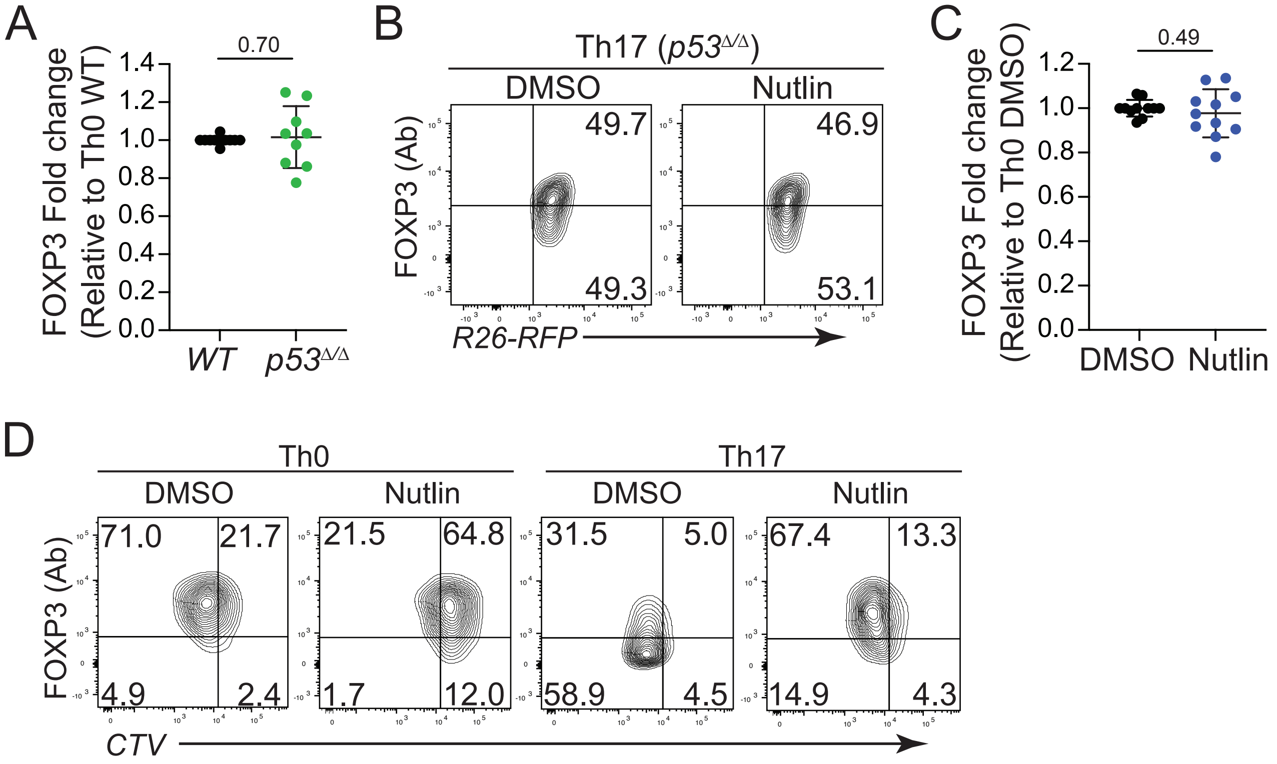


Supp. Figure 3 | p53 activity promotes the maintenance of Foxp3 expression. (A) Normalized FOXP3 expression of day 8 WT and p53 Δ/Δ Tregs in Th0 conditions. Data points represent biological replicates pooled from nine independent experiments. (B) Representative flow plots of day 8 FOXP3 expression in p53 Δ/Δ Tregs in Th17 conditions, treated with 10uM Nutlin or DMSO for 36 hours (starting day 3). (C) Normalized FOXP3 expression of day 8 WT Tregs in Th0 conditions, treated with 10 uM Nutlin or DMSO for 36 hours (starting day 3). Data points represent biological replicates pooled from ten independent experiments. (D) Representative flow plots of FOXP3 versus CellTrace proliferation dye for Tregs in Th0 or Th17 conditions, treated with 10uM Nutlin or DMSO for 36 hours (starting day 3). Tregs labeled with proliferation dye after washing out Nutlin on day 5, analysis on day 8. For all plots, *p<0.05, **p<0.01, ***p<0.001, ****p<0.0001 by unpaired Student's t-test, mean ± s.d.

Supp Figure 4: p53 is not necessary for Foxp3 induction in T cells

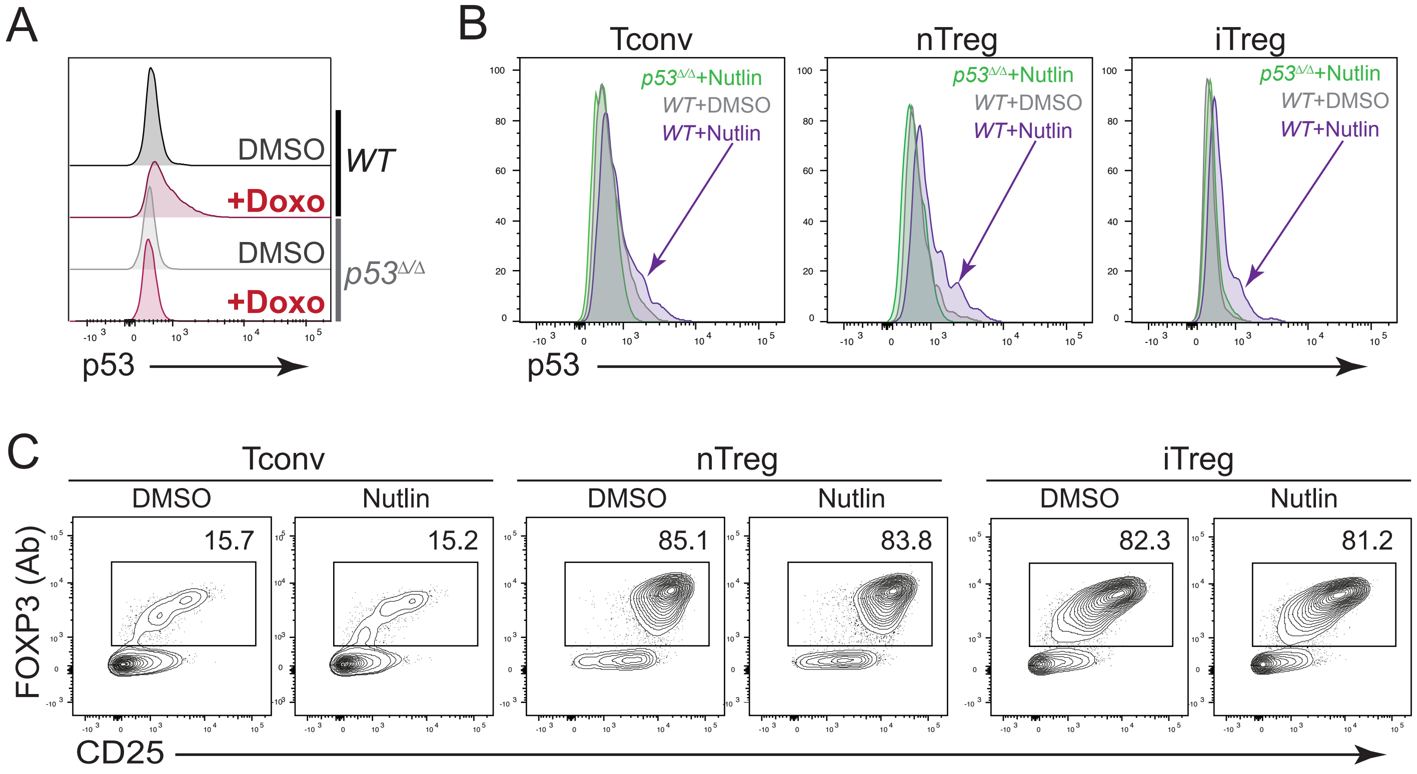


Supp. Figure 4 | p53 is not necessary for Foxp3 induction in T cells. (A) Representative histograms of p53 expression in 4 day in vitro-activated Tconv from WT or T.p53 Δ/Δ mice, treated with DMSO or 50 nM doxorubicin for 4 hours. (B) Representative histograms of p53 expression in Tconv, nTreg, or iTreg ChIP-qPCR samples treated with DMSO (WT) or 10 uM Nutlin (WT and p53 Δ/Δ) for 36 hours. (C) Representative flow plots of FOXP3 expression in WT Tconv, nTreg, or iTreg ChIP-qPCR samples treated with DMSO or 10 uM Nutlin for 36 hours. (D) ChIP-qPCR analysis of Cdkn1a in WT or p53 Δ/Δ Tconv, nTreg, or iTreg, treated with 10uM Nutlin or DMSO for 36 hours. Numbers indicate fold change relative to WT DMSO condition. Data points represent technical replicates, representative of at least two independent experiments.

Supp Figure 5: Treg-specific p53 deficiency leads to increased ex-Treg frequencies *in vivo* in Th17 cytokine-rich colon tissues

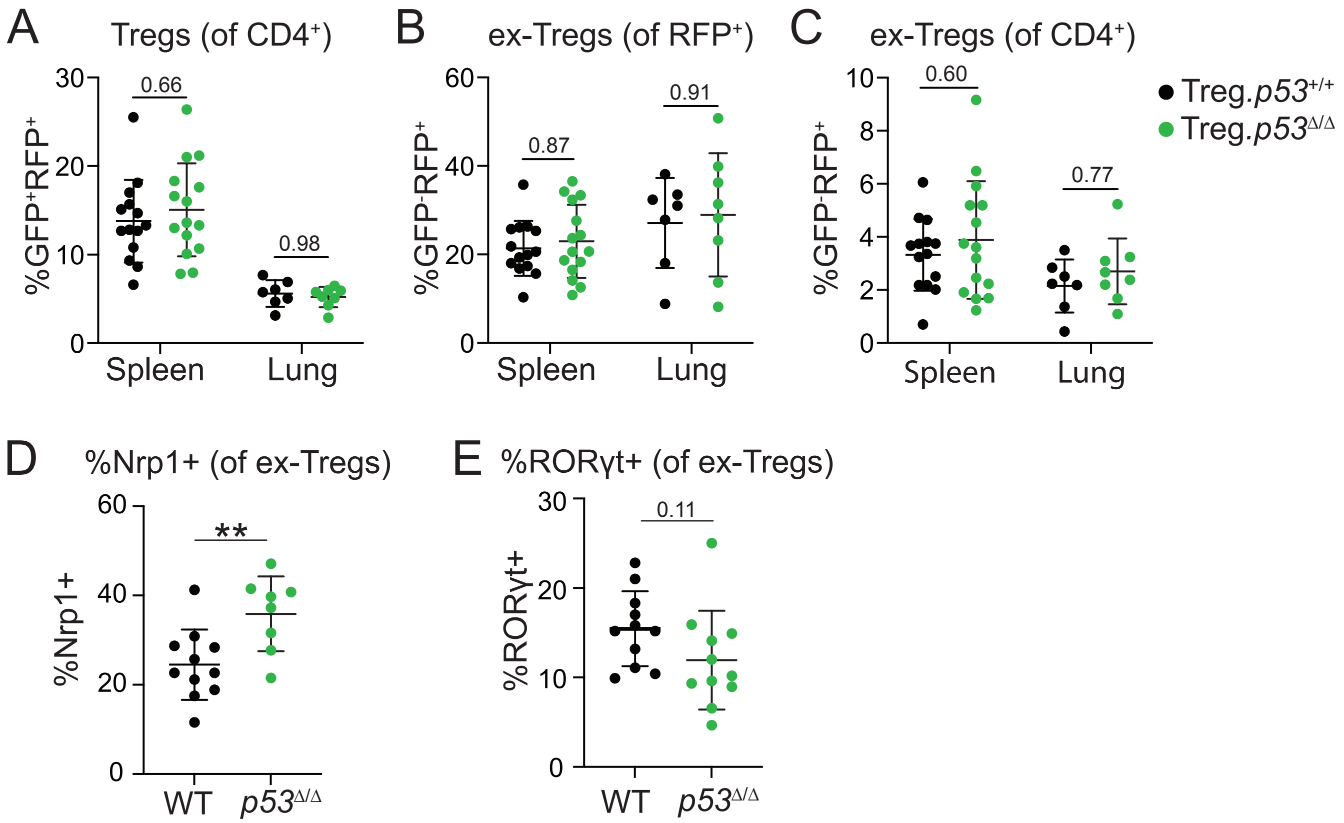


Supp. Figure 5 | Treg-specific p53 deficiency leads to increased ex-Treg frequencies in vivo in Th17 cytokine-rich colon tissues. (A) Frequency of GFP+RFP+ Tregs in the Spleen and Lung of WT and Treg.p53 Δ/Δ mice. Data from n=7-15 mice per genotype. (B-C) Frequency of ex-Tregs of RFP+ (B) or CD4+ (C) cells in the Spleen and Lung of WT and Treg.p53 Δ/Δ mice. Data from n=7-15 mice per genotype. (D-E) Frequency of Nrp1+ (D) and RORγt+ (E) cells among ex-Tregs in the colon of WT and Treg.p53 Δ/Δ mice. Data from n=8-11 mice per genotype. *p<0.05, **p<0.01, ***p<0.001, ****p<0.0001 by two-way ANOVA (A-C), or unpaired Student’s t-test (D-E), mean ± s.d.
